# Microtubule Binding Kinetics of Membrane-bound Kinesin Predicts High Motor Copy Numbers on Intracellular Cargo

**DOI:** 10.1101/627174

**Authors:** Rui Jiang, Steven Vandal, SooHyun Park, Sheereen Majd, Erkan Tüzel, William O. Hancock

**Affiliations:** Intercollege Program in Integrative and Biomedical Physiology, Pennsylvania State University, University Park, PA 16802; Department of Physics, Worcester Polytechnic Institute, Worcester, MA 01609; Department of Biomedical Engineering, Pennsylvania State University, University Park, PA 16802; Department of Biomedical Engineering, University of Houston, Houston, TX 77204

**Keywords:** Kinesin-1, lipid bilayers, kinetics, diffusion, long-range transport

## Abstract

Bidirectional vesicle transport along microtubules is necessary for cell viability and function, particularly in neurons. When multiple motors are attached to a vesicle, the distance a vesicle travels before dissociating is determined by the race between detachment of the bound motors and attachment of the unbound motors. Motor detachment rate constants (k_off_) can be measured via single-molecule experiments, but motor reattachment rate constants (k_on_) are generally unknown, as they involve diffusion through the bilayer, geometrical considerations of the motor tether length, and the intrinsic microtubule binding rate of the motor. To understand motor attachment dynamics during vesicle transport, we quantified the microtubule accumulation rate of fluorescently-labeled kinesin-1 motors in a 2D system where motors were linked to a supported lipid bilayer. From the first-order accumulation rate at varying motor densities, we extrapolated a k_off_ that matched single-molecule measurements, and measured a two-dimensional k_on_ for membrane-bound kinesin-1 motors binding to the microtubule. This k_on_ is consistent with kinesin-1 being able to reach roughly 20 tubulin subunits when attaching to a microtubule. By incorporating cholesterol to reduce membrane diffusivity, we demonstrate that this k_on_ is not limited by the motor diffusion rate, but instead is determined by the intrinsic motor binding rate. For intracellular vesicle trafficking, this two-dimensional k_on_ predicts that long-range transport of 100 nm diameter vesicles requires 35 kinesin-1 motors, suggesting that teamwork between different motor classes and motor clustering may play significant roles in long-range vesicle transport.

**Significance Statement:** Long-distance transport of membrane-coated vesicles involves coordination of multiple motors such that at least one motor is bound to the microtubule at all times. Microtubule attachment of a membrane-bound motor comprises two steps – diffusing through the lipid bilayer to a binding zone near the microtubule, followed by binding. Using a 2D supported lipid bilayer system, we show that membrane diffusion is not the limiting factor for motor attachment. This result suggests that in cells kinesin-1 binding kinetics are not altered by the membrane composition of vesicle cargos. The intrinsically slow binding properties of kinesin-1 suggest that divergent motor binding kinetics and motor clustering regulate long-range vesicle transport.

## Introduction

Survival and proper function of eukaryotic cells rely on the dynamics of numerous membranous organelles. Long-range transport of these cargos is carried out by the molecular motors kinesin and dynein, which walk towards the plus-(at cell periphery) and minus-ends (in perinuclear regions) of microtubules (MTs), respectively. Considerable work has been done to characterize the mechanochemical properties of kinesin and dynein by *in vitro* single-molecule studies (1–3). However, there is a significant knowledge gap between our understanding of *in vitro* motor biophysics and our understanding of *in vivo* vesicle transport. Generally, intracellular cargos have both kinesin and dynein simultaneously attached, and in a number of cases kinesins from multiple families are attached to the same cargo (4–6). Moreover, these motors are coupled through a lipid bilayer that surrounds the organelles, the effects of which have been studied only very recently (5, 7–10).

From a mechanical perspective, compared to attachment to a rigid surface, attachment to a lipid bilayer is expected to reduce motor performance because instead of moving the cargo along the microtubule, the motor can instead slip in the plane of the bilayer (7). On the other hand, being able to diffuse in the bilayer may enhance motor performance by allowing motors to diffuse to and accumulate on MTs. Geometry may also play a role; in a study of *in vitro* vesicle transport driven by myosin Va motors, fluid-phase membranes led to faster transport velocities than gel-phase membranes, an effect attributed to optimal centering of the vesicles on remaining motors following detachment of the trailing motor (8). Cellular studies have also revealed a role for membrane composition in regulating motor function. For instance, the kinesin-3 family member KIF16B exists in a monomeric state, but binding to phosphatidylinositol-(3) monophosphate (PI(3)P) on membranous cargos through its Phox homology (PX) domain converts KIF16B into superprocessive dimers that drive cargo transport (11). As another example, early phagosomes move on MTs bidirectionally, but as they mature, cholesterol is increasingly incorporated into the membrane, which results in clustering of dyneins into microdomains. This clustering allows dyneins to work cooperatively to generate greater minus-end forces, resulting in increasingly minus-end directed movements of late phagosomes (5).

Despite growing appreciation of the role of lipid membranes in regulating cargo transport, the mechanisms by which long-range vesicle transport is achieved are not well understood. The average travel distance of a vesicle depends on the average number of engaged motors, which in turn is determined by a race between motor attachment and detachment rates (Fig. 1A). Motor detachment is described by the first-order unbinding rate constant k_off_, which can be experimentally determined by dividing the single-molecule velocity by the run length (12–14). Measuring the motor attachment rate constant is more complicated because it involves two processes – diffusion of motors to a zone in which they are able to bind to the MT (described as a rate constant k_diffusion_ into a binding zone with area A_zone_), followed by MT binding, described by the first-order binding rate constant, k_attach_. Of these three parameters, k_attach_ for kinesin-1 has been measured in previous studies (10, 12), and bounds can be put on k_diffusion_ based on the diffusion constant of motors in the bilayer. In contrast, the area of the zone in which motors can bind to the MT is less well constrained, though the motor’s contour length provides an upper limit. How these parameters interact and whether cells modulate any or all of these parameters to regulate the speed and directionality of bidirectional transport are not understood. For instance, because different intracellular cargos have different lipid compositions (15–17), it is possible that the different diffusion constants alter motor binding kinetics and thus vesicle transport dynamics. This idea has not been experimentally tested, however.

**Fig. 1.**
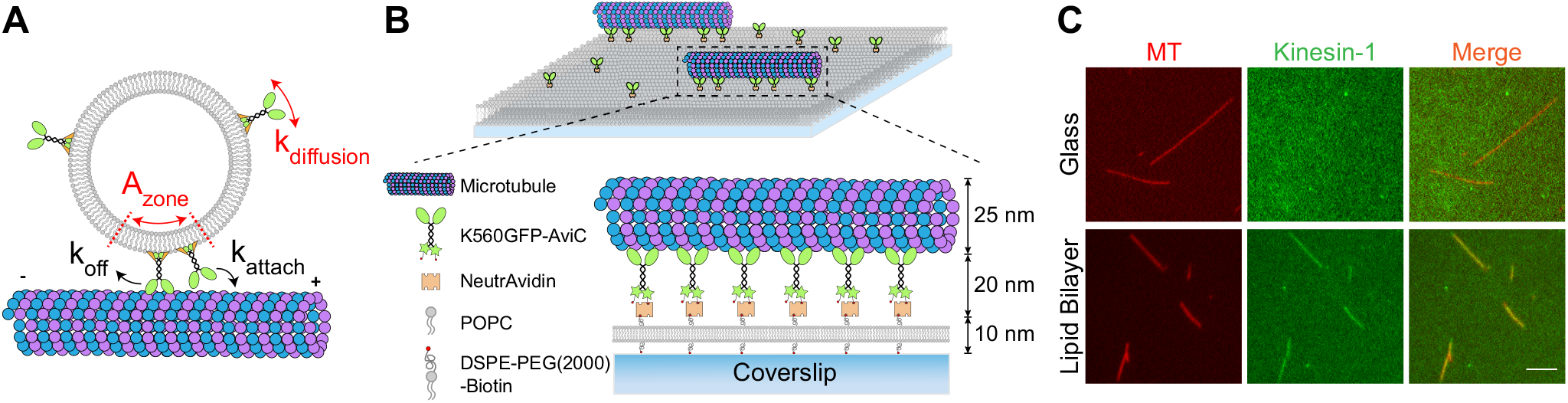
Visualization of membrane-mediated motor re-organization in a 2D supported lipid bilayer system. (A) Key kinetic parameters that determine average vesicle travel distance include the motor diffusion rate constant to the MT (k_diffusion_), the motor attachment rate constant (k_attach_), and the motor unbinding rate constant (k_off_). (B) Schematic illustrating linkage of biotinylated GFP-kinesin-1 (K560GFP-AviC) to DSPE-PEG(2000)-Biotin within a POPC lipid bilayer through NeutrAvidin. (C) Accumulation of membrane-bound kinesin-1 on MTs. Strong co-localization between kinesin-1 (green) and MTs (red) is seen only when kinesin-1 motors are bound to a lipid bilayer, not when motors are immobilized on a glass coverslip. Scale bar: 10 μm.

To better understand the principles that govern cargo transport by vesicle-bound motors, we used a 2D lipid bilayer system to approximate the local membrane environment where motors attach to a MT (Fig. 1B). By measuring the rate constant of accumulation of membrane-bound kinesin-1 on MTs, we obtained a two-dimensional bi-molecular rate constant for motor attachment, k_on_^2D^, for kinesin-1. By incorporating cholesterol to decrease the membrane diffusion constant, we reveal that this k_on_^2D^ is not limited by membrane diffusion but rather by the slow inherent binding rate of the motors. Using an analytical model, we find that this k_on_^2D^ can be explained by a small A_zone_ that is roughly the surface area of 20 tubulin dimers. Based on these findings, we conclude that enhanced long-distance transport of intracellular vesicles can be achieved by faster motor binding rates, slower motor unbinding rates, and by clustering of motors in membrane subdomains.

## Results and Discussion

### Membrane-bound Kinesin-1 Motors Accumulate on Microtubules

To measure the interrelationship between membrane diffusional kinetics and motor binding kinetics, we visualized GFP-labeled motors diffusing in a supported lipid bilayer system by total internal reflection fluorescence (TIRF) microscopy (Fig. 1B). This 2D system closely resembles the interface between a MT and a large membranous cargo, where the local membrane curvature is negligible. A lipid bilayer composed of POPC (1-palmitoyl-2-oleoyl-glycero-3-phosphocholine) spiked with small amount of DSPE-PEG(2000)-Biotin (1,2-distearoyl-*sn*-glycero-3-phosphoethanolamine-N-[biotinyl(polyethylene glycol)-2000]) was deposited on a cleaned glass coverslip, and biotinylated GFP-labeled *Drosophila* kinesin-1 motors (K560GFP-AviC (13)) were linked to DSPE-PEG(2000)-Biotin through NeutrAvidin. MTs were introduced into the chamber and changes in kinesin-1 distribution upon MT landing were monitored by local increases in fluorescence from the GFP tag. The fluid lipid bilayer enabled motors to freely diffuse within the membrane, resulting in strong accumulation of kinesin-1 on MTs (Fig. 1C). In a control experiment using kinesin-1 immobilized on a glass surface, no co-localization between motors and MTs was seen.

### Determining Motor Binding Kinetics from Accumulation Dynamics

To gain more insight into the kinetics of motor accumulation on the MT, we monitored the time course of motor fluorescence increase upon landing of a MT on the surface. Initially, motors were homogeneously distributed on the lipid bilayer, but upon MT landing, motors rapidly accumulated on MTs, reaching steady-state within seconds in the presence of 2 mM ATP (Movie S1, Fig. 2A, *SI Appendix*, Fig. S1). Consistent with a previous study (7), we observed MT gliding upon landing on the lawn of membrane-bound motors (*SI Appendix*, Fig. S1). To quantify motor accumulation, the integrated motor fluorescence in the region of the MT was fit to a single exponential function (Fig. 2C). We compared this motor accumulation rate constant, k_acc_, across different motor densities, and found that k_acc_ scaled linearly with motor density on the membrane (Movie S1, Fig. 2A, Fig. 2C, blue and green curves, Fig. 2D, blue curve). k_acc_ varied from 0.70 ± 0.03 s^−1^ (mean ± SEM, N = 3) at low motor density (~ 4 motors/μm^2^) up to 1.61 ± 0.12 s^−1^ (mean ± SEM, N = 35) at the highest motor density (~ 250 motors/μm^2^).

**Fig. 2.**
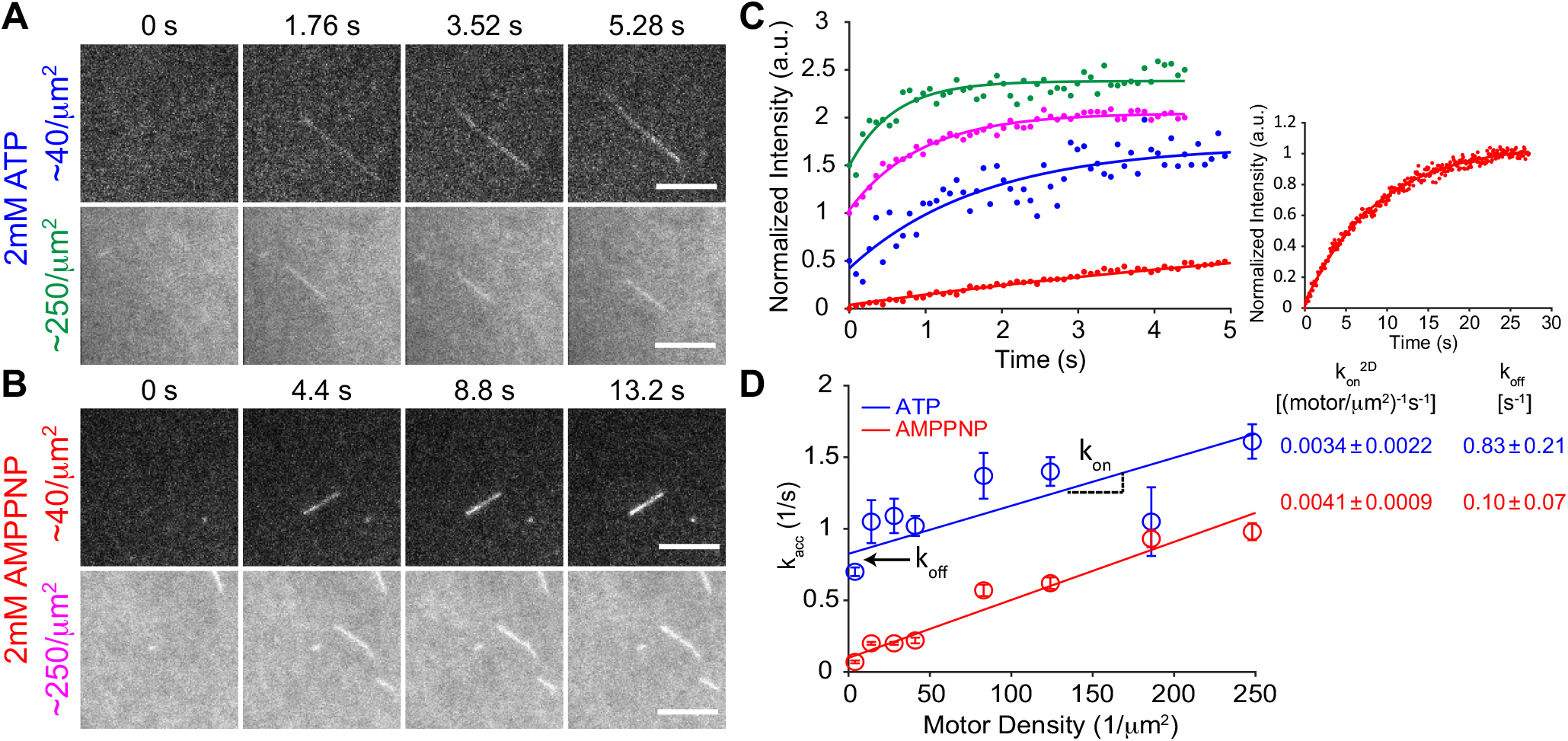
Quantification of k_on_^2D^ from the accumulation process. (A,B) Time-lapse snapshots of motor accumulation in 2 mM ATP (A) or 2 mM AMPPNP (B) at low (~40 motors/μm^2^) or high (~250 motors/μm^2^) motor densities. Upon MT landing at 0 s, the initially homogeneously distributed motors accumulate on the MT. Scale bar: 5 μm. See also *SI Appendix*, Fig. S1. (C) Time courses of integrated GFP intensity on MTs, with single exponential fit, color-coded for examples shown in (A,B). Inset: The entire curve (30 s) for the example in 2 mM AMPPNP at ~40 motors/μm^2^. Raw intensity is normalized and offset by 0.5 a.u. along the y-axis for visualization. (D) Accumulation rate (k_acc_, from exponential fits in C) versus kinesin-1 density in 2 mM ATP (blue) or 2 mM AMPPNP (red). Each data point represents mean of 3-41 measurements. Error bars indicate SEM.

To understand k_acc_ in terms of motor binding and unbinding rate constants, we applied a simple kinetic model of membrane-bound kinesin motors (Kin) binding to the surface-bound MT, as follows:

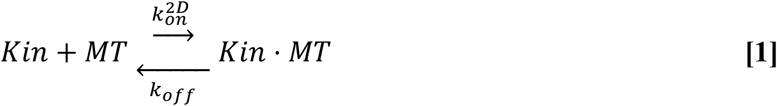

Here, k_on_^2D^ is the two-dimensional bimolecular association rate constant and k_off_ is the first-order motor dissociation rate constant. This system is identical to a reversible bimolecular interaction in solution (18), with the difference that instead of units M^−1^s^−1^, k_on_^2D^ has units (motors/μm^2^)^−1^s^−1^. It can be shown (see *SI Appendix, Supplementary Information Text* for details on derivation) that:

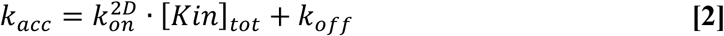

where [Kin]_tot_ is the surface density of kinesin-1. Thus, for the curves in Fig. 2D, a linear regression of k_acc_ as a function of motor density yields a line with a slope equal to the bimolecular association rate constant, k_on_^2D^, and a y-intercept equal to the first-order motor unbinding rate constant, k_off_. The fitted k_off_ of 0.83 ± 0.21 s^−1^ (mean ± 95% confidence bound of fit) was in good agreement with previous single-molecule measurements for these motors in saturating ATP, where k_off_ = velocity/run length (14, 19). From the linear fit, k_on_^2D^ in saturating ATP was 0.0034 ± 0.0022 (motor/μm^2^)^−1^s^−1^ (mean ± 95% confidence bound of fit).

One prediction of this model is that if we change the k_off_ without changing the k_on_^2D^, then the slope should be unaffected, and only the y-intercept should change. To test this prediction, we measured k_acc_ in the presence of 2 mM AMPPNP (Movie S1, Fig. 2B, Fig. 2C, red and magenta curves), instead of ATP. This non-hydrolyzable ATP analog prevents kinesin-1 dissociation from MT, thus reducing k_off_ to 0 s^−1^. Because ADP release is rate-limiting for kinesin in solution, the bimolecular interaction determining the k_on_^2D^ should be identical for the two nucleotides. As shown in the red curve in Fig. 2D, k_acc_ scaled linearly with motor density in AMPPNP, k_off_ was 0.10 ± 0.07 s^−1^ as expected, and the k_on_^2D^ of 0.0041 ± 0.0009 (motor/μm^2^)^−1^s^−1^ was similar to that in ATP. Thus, we have directly measured for the first time to our knowledge, the bimolecular association rate constant for membrane-bound motors binding to MTs. As shown below, this parameter allows one to predict the travel distance of a vesicle with known size, motor number (from which motor density can be calculated), and motor k_off_.

### Motor Attachment is Not Limited by Membrane Diffusion

The accumulation of membrane-bound motors on the MT can be considered as two sequential steps – motor diffusion (k_diffusion_) into a binding zone near the MT (A_zone_), followed by motor attachment to MT (k_attach_) from this binding zone (Fig. 1A). Depending on the diffusion constant and distance covered, k_on_^2D^ could in principle be determined by either the motor diffusion rate or the motor association rate when near the MT. Because different vesicle populations are known to have different lipid compositions (16, 17, 20), motor diffusivity could be a potential regulator of motor activity in cells.

To investigate whether diffusion or binding is rate-limiting for motor accumulation, we decreased the motor diffusion constant by incorporating 30 mol% cholesterol (the maximum amount under our conditions without membrane phase separation) into the POPC lipid bilayer. Fluorescence recovery after photobleaching (FRAP) measurements indicated that cholesterol incorporation resulted in a 3.8-fold decrease in the lipid diffusion coefficient from 0.95 ± 0.14 μm^2^/s for control to 0.25 ± 0.03 μm^2^/s with cholesterol (mean ± SD, N = 6 bilayers for each; Fig. 3A, *SI Appendix*, Table S1, Movie S2). To ensure that the FRAP measurements accurately reflected the diffusivity of motors in the membrane across different motor densities, we measured motor diffusion coefficients by single-molecule tracking in spiking experiments where only a small fraction of motors on the bilayer were labeled with GFP (*SI Appendix*, Fig. S2, Table S1). For the control POPC membrane, the single-molecule diffusion constant was 0.93 ± 0.03 μm^2^/s at ~40 motors/μm^2^ and 0.87 ± 0.02 μm^2^/s at ~250 motors/μm^2^ (mean ± 95% confidence bound of MSD fitting; N = 193 and 271, respectively). Thus, the single-molecule diffusion constants are in good agreement with the FRAP results and do not vary with motor density. Addition of 30% cholesterol decreased D to 0.46 ± 0.01 μm^2^/s at ~60 motors/μm^2^ and 0.53 ± 0.01 μm^2^/s at ~340 motors/μm^2^ (N = 301 and 254, respectively). This two-fold decrease measured by single-molecule tracking was less than the ~four-fold decrease measured by FRAP. In similar single-particle tracking experiments it was shown that single-motor diffusion coefficients have a broad distribution (21, 22). Because our measurement technique involved pre-bleaching a region of interest and tracking motors diffusing into the bleached area, we hypothesize that the diffusion constant in the cholesterol experiments preferentially selects for faster diffusing motors and hence may overestimate the mean single-molecule diffusion constant. Nonetheless, our measurements confirmed that incorporating cholesterol into the lipid bilayers effectively reduced motor diffusivity.

**Fig. 3.**
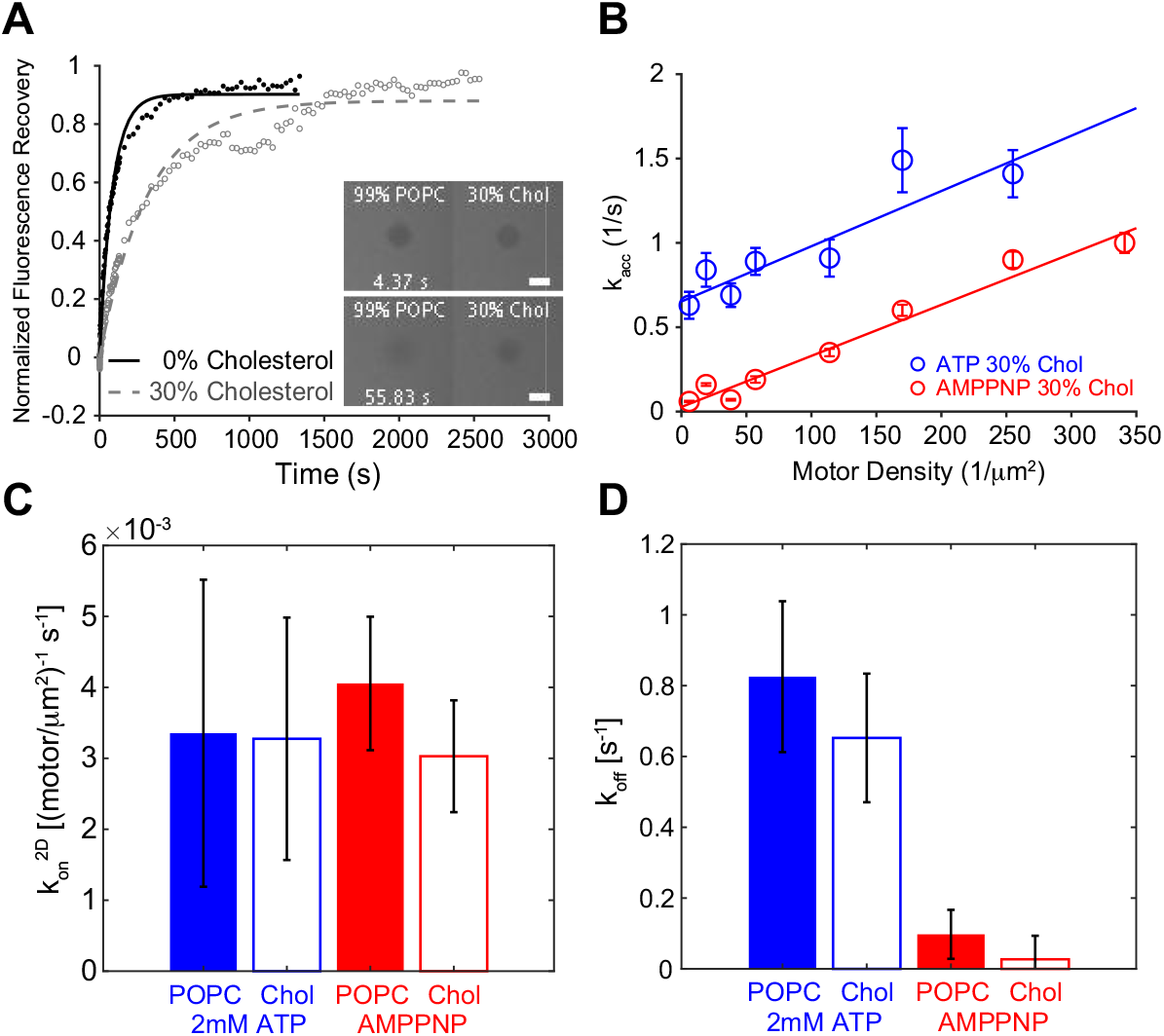
k_on_^2D^ of membrane-bound motors is not controlled by membrane diffusion. (A) Example traces of fluorescence recovery after photobleaching (FRAP) measurement of lipid bilayers without (black) or with 30% cholesterol (gray). Diffusion coefficients were calculated as 0.96 μm^2^/s and 0.25 μm^2^/s, respectively, from single exponential fitting. Scale bar: 30 μm. See *SI Appendix*, Fig. S2 for single-molecule diffusion estimates. (B) Motor accumulation rate versus kinesin-1 density in 2 mM ATP (blue) or 2 mM AMPPNP (red) in the presence of 30% cholesterol in the membrane. Each data point represents mean of 8-83 measurements. Error bars indicate SEM. (C,D) Summary of k_on_^2D^ (C) and k_off_ (D) of membrane-bound kinesin-1 motors under different membrane compositions and nucleotide conditions. Error bars indicate 95% confidence bound of fit.

We next quantified motor accumulation rate as a function of motor density in 2 mM ATP or AMPPNP on lipid bilayers containing 30% cholesterol (Fig. 3B). Addition of cholesterol into the membrane did not alter the motor attachment rate constant (k_on_^2D^ *=* 0.0033 ± 0.0017 (motors/μm^2^)^−1^s^−1^ in ATP, and 0.0030 ± 0.0008 (motors/μm^2^)^−1^s^−1^ in AMPPNP, Fig. 3C, 3D). This result seems surprising, but on closer inspection makes sense, as follows. If we assume that the zone from which motors can bind to the MT extends 50 nm on either side of the MT (roughly the kinesin tether length (23–25)), then from the 1D diffusion equation x^2^ = 2Dt, it should take motors an average of 1.3 ms (control) or 3-5 ms (plus cholesterol) to diffuse into and out of the zone. These times convert to rate constants orders of magnitude faster than 0.8 s^−1^ kinesin-1 unbinding rate constant, meaning that following detachment (or equivalently following attachment), motor populations inside and outside the binding zone will rapidly equilibrate. We therefore conclude that under our conditions, motor attachment to the MT is not limited by membrane diffusion. This suggests that kinesin-1 MT binding kinetics remain the same, regardless of membrane fluidity/composition of cargos, unless they are organized into higher-order structures.

### Measured k_on_^2D^ Predicts Kinesin-1 Can Explore 20 Tubulin Dimers During Attachment

To better understand our observed motor accumulation kinetics and to relate them to the physiological case of spherical vesicles of varying diameters, we developed an analytical model of the motor accumulation process. Because the motor diffusion rate within the membrane is considerably faster than motor binding kinetics, we made the assumption that the pool of motors within the binding zone is in rapid equilibrium with the pool of motors outside the binding zone. From our measured parameters k_on_^2D^ and σ, the 2D bimolecular on-rate and the motor surface density, respectively, the flux of motors binding to the MT (in motors/s) is equal to k_on_^2D^⋅σ (equivalent to k_on_ ⋅[motor] in solution). If we consider only those motors within the binding zone of a given tubulin subunit, then we can calculate an equivalent motor flux onto the MT as the number of motors in the zone, N, multiplied by a first-order motor attachment rate constant, k_attach_. In this case, the number of motors in the zone, N, is equal to σ⋅A_zone_, where σ is the motor density and A_zone_ is the area. Uniting these two representations of the motor flux onto the MT,

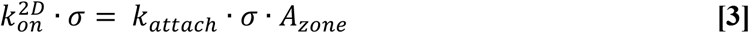

Thus,

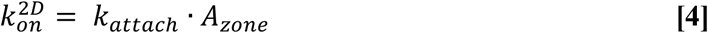

Our experimentally determined k_on_^2D^ was 0.0034 (motor/μm^2^)^−1^s^−1^. The first-order kinesin attachment rate constant, k_attach_ has been previously calculated from a lipid vesicle system to be 4.7 s^−1^ (10), from a DNA scaffold system to be 4.6 s^−1^ (12), and from a bead system to be 5 s^−1^ (26). Although the 2D lipid bilayer system is a third geometry, we make the tentative assumption that the first-order association rate constant also applies here. Thus, we can calculate an estimate for the area around a given tubulin subunit from which motors can bind as A_zone_ = 680 nm^2^. This value is considerably smaller than the ~8000 nm^2^ area predicted by a model in which all of the motors within a 50 nm motor contour length can bind to a given tubulin, an approach that was taken by others (5, 27). If we assume that the motor reach is the determining factor, then the area of motors accessible to a given tubulin subunit should be equivalent to the area of tubulin subunits accessible to a given motor. Thus, we can interpret the A_zone_ of 680 nm^2^ as kinesin-1 being able to stretch ~15 nm (a circle with 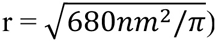), or equivalently, being able to explore ~20 tubulin subunits (680 nm^2^ /(8 nm × 4 nm)). This reach is far from the motor’s contour length of 80 nm, but close to the distance that kinesin-1 holds its cargo from the MT, previously measured to be ~17 nm (23, 28).

### Slow Attachment Predicts High Motor Densities for Long-range Vesicle Transport

Achieving run lengths greater than the single-motor distance requires multiple motors on a cargo simultaneously binding to the MT such that when one motor dissociates, others are there to continue transport. Using the kinetic parameters measured here, we carried out simulations to predict the number of kinesin-1 motors required to achieve long-range transport of vesicles of varying diameters. The vesicle motor density, σ, was defined as the number of motors on the vesicle divided by the membrane surface area of the vesicle. The flux of vesicle-bound motors binding to the MT was defined as k_on_2D_·σ, and the flux of motors detaching from the MT was determined by k_off_ times the number of MT-bound motors. A naïve prediction of this system is that for vesicles smaller than the motor contour length, every motor can access the MT, so having multiple motors on a vesicle leads to longer run lengths. Consistent with this prediction, for vesicles of 30 nm in diameter, 2 motors are sufficient to double the run length (Fig. 4A), and 6 motors can carry the vesicle for 10 μm on average. However, analogous to the slow accumulation in the supported lipid bilayer system, we found that for larger cargos, achieving multi-micron travel distances requires higher motor numbers. For instance, a 100 nm vesicle needed ~35 kinesin-1 motors to travel 10 μm, and a 500 nm vesicle required 800 motors to travel the same distance (Fig. 4A).

**Fig. 4.**
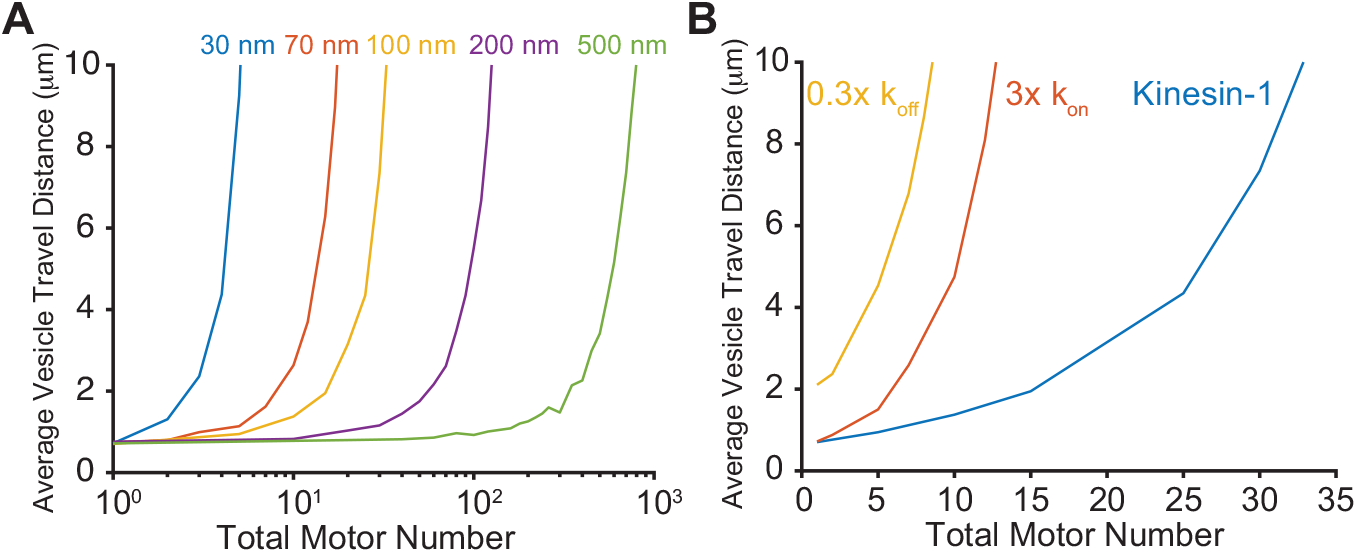
Slow motor attachment rate constant predicts high motor density for long-range vesicle transport. (A) Simulation of vesicle travel distance versus motor number per vesicle for vesicles of varying diameters. (B) Simulation of travel distance versus motor number for a 100 nm vesicle, using a different motor binding rate constant or a different unbinding rate constant.

Although this requirement for high motor numbers per vesicle seems surprising, it is in fact consistent with a number of published optical trap experiments, where single-molecule activity is routinely achieved by incubating ~500-1000 nm diameter beads with a thousand-fold excess of motors (29, 30). For vesicles larger than the motor contour length, motor-MT interactions are naturally geometrically constrained; for instance it has been calculated that for a 760 nm diameter vesicle, only 4% of dynein randomly distributed on the surface can reach the MT, assuming a dynein contour length of 70 nm (5). Our conclusion that the binding zone for a motor is much less than its contour length further shifts this percentage down and provides a kinetic description of the process. In published work, smaller cargos were also found to have a similar requirement of high kinesin-1 copy numbers for long-distance transport. Using 100 nm diameter beads, Beeg *et al.* found that the run length only marginally increased when up to 73 motors were bound to the bead (26), whereas we predict 35 motors are sufficient to carry a 100 nm vesicle for 10 μm on average (Fig. 4A). This difference suggests that motor diffusion in a lipid bilayer may facilitate transport. One plausible explanation for this is that when motors are bound to a rigid cargo, only a small number of motors are present in the binding zone, thus motors binding to the MT result in motor depletion in the zone. In contrast, if motors are bound to a lipid vesicle, motor depletion is negligible because the motor population inside the zone quickly equilibrates with the pool outside the zone through diffusion.

The motor copy numbers that we predict based on our accumulation measurements are somewhat higher than previous measurements using purified neuronal vesicles and phagosomes. Using quantitative Western blots or photobleaching, it was estimated that ~90 nm vesicles contain on average 3-10 dynein, 1-3 kinesin-1, and 2-6 kinesin-2 motors (6, 31). One hypothesis to account for these low motor numbers is that motors from different families have different binding kinetics that could improve their transport efficiency in teams and thereby reduce the number of motors required for long-range transport. For instance, kinesin-2 has been shown to have a 4-fold faster k_on_ compared to kinesin-1, suggesting it may play a tethering role to maximize attachment of a vesicle to the MT (12). Also, activated mammalian dynein has a k_off_ on the order of 0.04 s^−1^ (32), meaning that slowly detaching dyneins can hold a vesicle near the MT, enabling kinesins that detach to rapidly rebind and continue bidirectional transport. To test this hypothesis, we ran simulations where we varied k_on_ or k_off_, and found that transport efficiency increased drastically with either a faster k_on_ or slower k_off_ (Fig. 4B). If the motor had a 3-fold faster k_on_, then instead of 35 motors, only 13 motors were needed to carry a 100 nm vesicle for 10 μm. A 3-fold slower k_off_ had an even more prominent effect – only 9 motors were sufficient to carry the vesicle for the same distance. Motor clustering into membrane subdomains is another factor that could enhance long-distance vesicle transport. For instance, in late phagosomes dynein has been shown to cluster into microdomains (5), which is expected to increase its k_on_ dramatically because motors no longer diffuse in the membrane and their clustering significantly increases the local motor density. In analogy to motor clustering within lipid microdomains, motors closely packed on DNA scaffolds generally have long run lengths with low motor copy numbers. Two kinesin-1 motors spaced 15 or 23 nm apart on a DNA scaffold doubled the run length (12, 33), and four kinesin-1 spaced 23 nm apart (33) or seven kinesin-1 spaced 28 nm apart (34) exhibited a 7-fold longer run lengths.

### Conclusions

Organelles and other membranous cargos need to travel tens of μm or longer to reach their cellular destinations, which is achieved by teamwork in a group of motors that only travel several μm individually. The distance that a cargo can travel is ultimately determined by the race between attachment of the free motors and detachment of the bound motors. In this study, we dissected four key parameters in this kinetic process. From the motor accumulation on a lipid bilayer, we measured k_off_, which matches with previous single-molecule studies, and the two-dimensional bimolecular association rate constant, k_on_^2D^. This on-rate parameter encompasses: i) the membrane diffusion rate constant k_diffusion_, ii) the binding zone A_zone_ within which motors can access the MT, and iii) k_attach_, the first-order attachment rate constant when motors are within the binding zone. k_on_^2D^ remained unchanged when membrane diffusivity was reduced by cholesterol, indicating that motor attachment is not limited by membrane diffusion and suggesting that changes in membrane fluidity are not an effective mechanism by which cells can regulate transport properties of different vesicle cargos by kinesin-1. Using the reported value of k_attach_, our theoretical work suggests an A_zone_ smaller than that predicted by the motor contour length, and one that is close to the area of ~20 tubulin dimers. This small zone of interaction explains the intrinsically slow binding of kinesin-1 to the MT and the prediction that relatively high motor densities are required for long-range vesicle transport. This work suggests teamwork between dissimilar motors can enhance vesicle transport, and leaves open the possibility that some vesicles achieve long-distance transport by clustering motors in membrane microdomains or through scaffold proteins. As suggested from recent work on dynein (5), this clustering may be a mechanism to regulate vesicle moving directionality during bidirectional transport in cells.

## Materials and Methods

### Protein Constructs and Purification

The K560GFP-AviC construct was made by inserting the biotin ligase consensus sequence (AviTag) (13) between the eGFP tag and the 6x His tag in a previously described K560GFP construct (12). Insertions were carried out using Q5 site-directed mutagenesis (New England Biosciences). Motors were expressed polycistronically with BirA biotin ligase, in BL21 (DE3) bacteria (New England Biosciences) and purified by affinity chromatography, as previously described (12, 13, 35). Motor concentration was quantified by GFP absorbance or fluorescence at 488 nm.

### Formation of Supported Lipid Bilayers (SLBs)

1-palmitoyl-2-oleoyl-glycero-3-phosphocholine (POPC), 1,2-disteroyl-*sn*-glycero-3-phosphoethanolamine-N-[biotinyl(polyethylene glycol)-2000] (DSPE-PEG(2000)-Biotin) and cholesterol were purchased from Avanti. Lipids were dissolved in chloroform and mixed in desired molar ratio. 1,2-dioleoyl-*sn*-glycero-3-phosphoethanolamine labeled with Atto 647N (Atto 647N DOPE) (Sigma) was incorporated into all lipid bilayers for visualization. Lipid composition without cholesterol was POPC:DSPE-PEG(2000) Biotin:Atto 647N DOPE at 99:0.1-0.6:0.05, and the composition with cholesterol was POPC:cholesterol:DSPE-PEG(2000) Biotin:Atto 647N DOPE at 69:30:0.1-0.6:0.05. The bulk of chloroform in the lipid mixture was evaporated in a rotary evaporator for 1.5 h. Lipids were left under vacuum for another 3 h to remove any trace chloroform. The dry lipid film was rehydrated in BRB80 (80 mM Pipes, 1 mM EGTA, 1 mM MgCl_2_; pH 6.9) to a final lipid concentration of 0.5 mg/ml, and the lipid suspension was subjected to 10 freeze-thaw cycles in liquid nitrogen and warm water bath. Small unilamellar vesicles (SUVs) were formed by passing the lipid solution 11 times through a 100 nm polycarbonate membrane, followed by 21 times through a 30 nm membrane, using a mini-extruder (Avanti).

Glass coverslips (#1.5, 24×30 mm, Corning) were cleaned as previously described (36). Coverslips were boiled in 7x detergent (MP Biomedicals) diluted 1:7 with 18.2 MΩ water for 2 h. Coverslips were then rinsed thoroughly with 18.2 MΩ water, blown dry with nitrogen, and annealed at 550 °C for 6 h. Coverslips were cleaned by a plasma cleaner (Harrick Plasma) immediately before formation of SLBs. Polydimethylsiloxane (PDMS) spacers were made using Sylgard 184 silicone elastomer kit (Dow Corning). Cured PDMS was cut by a biopsy punch (Acuderm) to make spacers 6 mm inner diameter and ~2 mm thick that were adhered to clean coverslips. 50 μl SUVs was added to the well and incubated for 10 min, followed by thorough wash in BRB80 to remove excess SUVs.

### Accumulation Assay

Taxol-stabilized microtubules (MTs) were polymerized from bovine brain tubulin, as described previously (12–14, 35). Rhodamine-labeled MTs (rhodamine:unlabeled = 1:3) were used in experiments in Fig. 1C, unlabeled MTs were used otherwise. MTs were sheared by passing twice through a 30g needle. Motor-functionalized bilayers were formed by binding pre-formed kinesin-NeutrAvidin complexes to biotinylated lipid bilayers, as follows. 200 nM kinesin was incubated with 2 μM NeutrAvidin (Thermo Scientific) in BRB80 containing 100 μM AMPPNP, 10 μM taxol, and 1 mM DTT for 10 min at room temperature (RT). Excess of NeutrAvidin ensured that all biotinylated motors were occupied, without cross-linking of motors. To remove excess NeutrAvidin, the solution was incubated 10 min with 1 μM MT, centrifuged for 10 min in an Airfuge (Beckman-Coulter), inactive motors and excess NeutrAvidin in the supernatant were discarded. The pellet containing MTs with bound kinesin-NeutrAvidin (K-NA) was resuspended in motor buffer (1 μM taxol, 2 mM ATP, 20 mM D-glucose, 1 mM DTT in BRB80), incubated for 10 min at RT to release active motors, followed by an Airfuge spin to remove MTs with irreversibly bound motors.

SLBs were equilibrated with motor buffer, 50 μl of K-NA solution was then introduced, followed by a 10 min incubation to allow motors to bind to the SLB. Excess motors were removed by a wash of motility buffer (1 μM taxol, 2 mM ATP or AMPPNP, 1 mM DTT, 40 mM D-glucose, 0.02 mg/ml glucose oxidase, 0.008 mg/ml catalase in BRB80). 16-32 nM MTs in motility buffer were introduced immediately before imaging, and accumulation of GFP-labeled motors was imaged using a Nikon TE2000 TIRF microscope (except for Fig. 1C, which was captured using an epifluorescence microscope) at 10 – 11.4 fps, as described previously (14).

### Motor Density Quantification and Single-Motor Diffusion Determination

At low motor densities, SLBs containing 0.1% DSPE-PEG(2000)-Biotin were incubated with NeutrAvidin spiked with varying amount of K-NA to control the motor density. We directly counted motor densities at spiking ratios of 1:500 to 1:50, giving motor densities of 41 /μm^2^ and 57 /μm^2^ for full motor occupancy for SLBs containing 0.1% DSPE-PEG(2000)-Biotin, without and with 30% cholesterol, respectively. To achieve high motor densities, we increased the amount of DSPE-PEG(2000)-Biotin in the SLBs (up to 0.6%) and incubated the SLBs with saturating levels of K-NA. We measured the mean GFP intensities at varying surface motor densities under constant TIRF illumination and exposure time. The fluorescence intensity scaled linearly with equivalent biotin concentration in the membrane (DSPE-PEG(2000)-Biotin (%) × spiking ratio, *SI Appendix*, Fig. S3). This allowed us to scale up the number for high motor densities.

To quantify the single-molecule motor diffusion constant, we attached K560AviC spiked with K560GFPAviC to the SLBs, as follows. SLBs were first incubated with NeutrAvidin spiked with K560GFPAviC-NA complexes. After removing the unbound proteins, SLBs were then incubated with saturating level of K560AviC to occupy all the NeutrAvidin sites. Motor diffusion was imaged by TIRF microscopy at 10 fps.

### Fluorescence Recovery After Photobleaching (FRAP)

To obtain sufficient signal-to-noise ratio for FRAP measurement, 0.5 mol% Atto 647N DOPE was incorporated into SLB samples. Images were captured using a Nikon Eclipse Ti microscope equipped with a Plan Apo 10x DIC N1 objective and an attached digital complementary metal-oxide semiconductor (CMOS) camera (ORCA-Flash 4.0 V2, Hamamatsu) in conjunction with NIS-Elements (Nikon) software. SLBs were excited with a 640 nm light source (Aura II, Lumencor). A 15 mW 561 nm laser (LU-N3 Laser Unit, Nikon) equipped with a Bruker Miniscanner was used to bleach Atto 647N DOPE at 50-80% output power for 3 s. The laser beam had an effectively cylindrical profile with a diameter of 30 μm. To minimize photobleaching during recovery, images were captured immediately after bleaching every 40-60 ms for 2 s, then every 500 ms for 10 s, followed by every 3 s for 2 min, and finally switched to every 30 s. Data analysis was carried out as previously described (36), using a correction factor of 1.

### Data Analysis

Accumulation image sequences were analyzed in Fiji (37). The trace that a newly landing MT glided along was tracked manually and the GFP intensity profile along the trace was plotted over time. The frame that the MT landed was defined by the slope maximum on a plot of average GFP intensity along the trace versus time. After background correction, the trace was fitted by a single exponential function from which k_acc_ was obtained. A weighted (by inverse of SEM) linear regression between k_acc_ and surface active motor density by linear least square fitting was carried out in MATLAB (The MathWorks).

Diffusion of single motors was tracked using FIESTA (38). Displacement data of all trajectories were calculated for discrete lag times, and the displacement data were cumulated to calculate the average MSD for each time point. To obtain motor diffusion coefficients, the first 10 points of the MSD were fitted with a line weighted by the inverse of standard errors.

### Simulation of Vesicle Transport

Vesicle transport was simulated by modifying a MATLAB code that was previously used to simulate single-molecule runs (35). The vesicle diameter, D, and total motor number, N, were defined at the beginning of the simulation. Each state was defined by the number of engaged motors, i. The transition rate from state i to state i+1 was defined as k_on_^2D^∙(N -i) / (π∙(D/2)^2^). The transition rate from state i back to state i-1 was defined as k_off_ ∙ i. Vesicle dwell times on the MT were simulated based on the transition matrix. Average vesicle transport distances were calculated by multiplying the simulated dwell time by the kinesin-1 velocity of 600 nm/s (13).

## Acknowledgements

We are grateful to David Arginteanu for his help with protein preparation, Paul Cremer and Simou Sun for advice on supported lipid bilayers, Codey Henderson for assistance on FRAP experiments, members from Hancock laboratory and Cremer laboratory for helpful discussions, and the reviewers for insightful comments.

This work was funded by National Institutes of Health (NIH) grant R01 GM121679 to W.O.H. and E.T.R.J. was supported by NIH T32 GM108563.

## Author Contributions

R.J., W.O.H. designed research; R.J., S.V., S.P. performed research; R.J., S.V., S.M., E.T. and W.O.H. analyzed data; and R.J. and W.O.H. wrote the paper.

## References

1. W. O. Hancock, The Kinesin-1 Chemomechanical Cycle: Stepping Toward a Consensus. Biophys J 110, 1216–1225 (2016).

2. D. A. Grotjahn et al., Cryo-electron tomography reveals that dynactin recruits a team of dyneins for processive motility. Nat Struct Mol Biol 25, 203–207 (2018).

3. L. Urnavicius et al., Cryo-EM shows how dynactin recruits two dyneins for faster movement. Nature 554, 202–206 (2018).

4. G. T. Shubeita et al., Consequences of motor copy number on the intracellular transport of kinesin-1-driven lipid droplets. Cell 135, 1098–1107 (2008).

5. A. Rai et al., Dynein Clusters into Lipid Microdomains on Phagosomes to Drive Rapid Transport toward Lysosomes. Cell 164, 722–734 (2016).

6. A. G. Hendricks et al., Motor coordination via a tug-of-war mechanism drives bidirectional vesicle transport. Curr Biol 20, 697–702 (2010).

7. R. Grover et al., Transport efficiency of membrane-anchored kinesin-1 motors depends on motor density and diffusivity. Proc Natl Acad Sci U S A, (2016).

8. S. R. Nelson, K. M. Trybus, D. M. Warshaw, Motor coupling through lipid membranes enhances transport velocities for ensembles of myosin Va. Proc Natl Acad Sci U S A 111, E3986–3995 (2014).

9. D. R. Klopfenstein, M. Tomishige, N. Stuurman, R. D. Vale, Role of phosphatidylinositol(4,5)bisphosphate organization in membrane transport by the Unc104 kinesin motor. Cell 109, 347–358 (2002).

10. C. Leduc et al., Cooperative extraction of membrane nanotubes by molecular motors. Proc Natl Acad Sci U S A 101, 17096–17101 (2004).

11. V. Soppina et al., Dimerization of mammalian kinesin-3 motors results in superprocessive motion. Proc Natl Acad Sci U S A 111, 5562–5567 (2014).

12. Q. Feng, K. J. Mickolajczyk, G. Y. Chen, W. O. Hancock, Motor Reattachment Kinetics Play a Dominant Role in Multimotor-Driven Cargo Transport. Biophys J 114, 400–409 (2018).

13. K. J. Mickolajczyk et al., Kinetics of nucleotide-dependent structural transitions in the kinesin-1 hydrolysis cycle. Proc Natl Acad Sci U S A 112, E7186–7193 (2015).

14. S. Shastry, W. O. Hancock, Neck linker length determines the degree of processivity in kinesin-1 and kinesin-2 motors. Curr Biol 20, 939–943 (2010).

15. P. Sanghavi et al., Coin Tossing Explains the Activity of Opposing Microtubule Motors on Phagosomes. Curr Biol, (2018).

16. N. Rocha et al., Cholesterol sensor ORP1L contacts the ER protein VAP to control Rab7-RILP-p150 Glued and late endosome positioning. J Cell Biol 185, 1209–1225 (2009).

17. G. van Meer, D. R. Voelker, G. W. Feigenson, Membrane lipids: where they are and how they behave. Nat Rev Mol Cell Biol 9, 112–124 (2008).

18. T. D. Pollard, E. M. De La Cruz, Take advantage of time in your experiments: a guide to simple, informative kinetics assays. Mol Biol Cell 24, 1103–1110 (2013).

19. K. J. Mickolajczyk, W. O. Hancock, Kinesin Processivity Is Determined by a Kinetic Race from a Vulnerable One-Head-Bound State. Biophys J 112, 2615–2623 (2017).

20. D. Pathak, R. Mallik, Lipid - Motor Interactions: Soap Opera or Symphony? Curr Opin Cell Biol, (2016).

21. S. Pyrpassopoulos et al., Force Generation by Membrane-Associated Myosin-I. Sci Rep 6, 25524 (2016).

22. L. B. Sagle et al., Single plasmonic nanoparticle tracking studies of solid supported bilayers with ganglioside lipids. J Am Chem Soc 134, 15832–15839 (2012).

23. J. Kerssemakers, J. Howard, H. Hess, S. Diez, The distance that kinesin-1 holds its cargo from the microtubule surface measured by fluorescence interference contrast microscopy. Proc Natl Acad Sci U S A 103, 15812–15817 (2006).

24. J. M. Scholey, J. Heuser, J. T. Yang, L. S. Goldstein, Identification of globular mechanochemical heads of kinesin. Nature 338, 355–357 (1989).

25. N. Hirokawa et al., Submolecular domains of bovine brain kinesin identified by electron microscopy and monoclonal antibody decoration. Cell 56, 867–878 (1989).

26. J. Beeg et al., Transport of beads by several kinesin motors. Biophys J 94, 532–541 (2008).

27. R. P. Erickson, Z. Jia, S. P. Gross, C. C. Yu, How molecular motors are arranged on a cargo is important for vesicular transport. PLoS Comput Biol 7, e1002032 (2011).

28. R. H. Miller, R. J. Lasek, Cross-bridges mediate anterograde and retrograde vesicle transport along microtubules in squid axoplasm. The Journal of cell biology 101, 2181–2193 (1985).

29. S. S. Rosenfeld, P. M. Fordyce, G. M. Jefferson, P. H. King, S. M. Block, Stepping and stretching. How kinesin uses internal strain to walk processively. J Biol Chem 278, 18550–18556 (2003).

30. Q. Li, S. J. King, A. Gopinathan, J. Xu, Quantitative Determination of the Probability of Multiple-Motor Transport in Bead-Based Assays. Biophys J 110, 2720–2728 (2016).

31. A. R. Chaudhary, F. Berger, C. L. Berger, A. G. Hendricks, Tau directs intracellular trafficking by regulating the forces exerted by kinesin and dynein teams. Traffic 19, 111–121 (2018).

32. R. J. McKenney, W. Huynh, M. E. Tanenbaum, G. Bhabha, R. D. Vale, Activation of cytoplasmic dynein motility by dynactin-cargo adapter complexes. Science 345, 337–341 (2014).

33. K. Furuta et al., Measuring collective transport by defined numbers of processive and nonprocessive kinesin motors. Proc Natl Acad Sci U S A 110, 501–506 (2013).

34. N. D. Derr et al., Tug-of-war in motor protein ensembles revealed with a programmable DNA origami scaffold. Science 338, 662–665 (2012).

35. G. Y. Chen, D. F. Arginteanu, W. O. Hancock, Processivity of the kinesin-2 KIF3A results from rear head gating and not front head gating. J Biol Chem 290, 10274–10294 (2015).

36. A. M. Sendecki, M. F. Poyton, A. J. Baxter, T. Yang, P. S. Cremer, Supported Lipid Bilayers with Phosphatidylethanolamine as the Major Component. Langmuir 33, 13423–13429 (2017).

37. J. Schindelin et al., Fiji: an open-source platform for biological-image analysis. Nature methods 9, 676 (2012).

38. F. Ruhnow, D. Zwicker, S. Diez, Tracking single particles and elongated filaments with nanometer precision. Biophys J 100, 2820–2828 (2011).

