## Supplemental Information for "Microtubule Binding Kinetics of Membrane-bound Kinesin Predicts High Motor Copy Numbers on Intracellular Cargo"

**This PDF file includes:**

Supplementary text
Figures S1 to S3
Table S1
Legends for Movies S1 to S2

**Other supplementary materials for this manuscript include the following:**

Movies S1 to S2

### Supplementary Information Text

#### Analytical model for motor accumulation rate

The motor accumulation rate in Fig. 2, $k_{acc} = k_{on}^{2D} \cdot [Kin]_{tot} + k_{off}$, was derived as follows. The reversible binding interaction of surface-bound kinesin to microtubules that land on the surface is:

$$Kin + MT \underset{k_{off}}{\overset{k_{on}^{2D}}{\rightleftarrows}} Kin \cdot MT \quad \textbf{[1]}$$

where [Kin] is the free kinesin concentration, [MT] is the concentration of available binding sites on the MT, and [Kin·MT] is the concentration of MT-bound kinesin. The rate of accumulation of kinesin on the microtubule can be expressed as:

$$\frac{d[Kin \cdot MT]}{dt} = k_{on}^{2D} \cdot [Kin] \cdot [MT] - k_{off} \cdot [Kin \cdot MT] \quad \textbf{[2]}$$

If the assumption is made that the amount of MT-bound kinesin-1, [Kin·MT], is negligible compared to the total pool of membrane-bound motors, $[Kin]_{tot}$ ($[Kin \cdot MT] << [Kin]_{tot}$), then $[Kin] = [Kin]_{tot}$ at all times. Because the total concentration of tubulin binding sites, $[MT]_{tot}$, is conserved, it follows that

$$[MT] = [MT]_{tot} - [Kin \cdot MT] \quad \textbf{[3]}$$

Plugging in and rearranging,

$$\frac{d[Kin \cdot MT]}{dt} = k_{on}^{2D} \cdot [Kin]_{tot} \cdot [MT]_{tot} - \left(k_{off} + k_{on}^{2D} \cdot [Kin]_{tot}\right) \cdot [Kin \cdot MT] \quad \textbf{[4]}$$

At steady-state, d[Kin·MT]/dt = 0, thus

$$k_{on}^{2D} \cdot [Kin]_{tot} \cdot [MT]_{tot} - \left(k_{off} + k_{on}^{2D} \cdot [Kin]_{tot}\right) \cdot [Kin \cdot MT]_{ss} = 0 \quad \textbf{[5]}$$

Where $[Kin \cdot MT]_{ss}$ is the steady-state concentration of MT-bound kinesin motors. The equation can be rearranged to solve for the steady-state motor accumulation:

$$[Kin \cdot MT]_{ss} = \frac{k_{on}^{2D} \cdot [Kin]_{tot}}{k_{off} + k_{on}^{2D} \cdot [Kin]_{tot}} [MT]_{tot} \quad \textbf{[6]}$$

Using the boundary conditions [Kin·MT] = 0 at t = 0 and $[Kin \cdot MT] = [Kin \cdot MT]_{ss}$ at t = ∞, Eq. **4** can be solved for the timecourse of motors bound to the MT, [Kin·MT]:

$$[Kin \cdot MT] = \frac{[MT]_{tot}}{1 + k_{off}/(k_{on}^{2D} \cdot [Kin]_{tot})} \left(1 - e^{-(k_{on}^{2D} \cdot [Kin]_{tot} + k_{off}) \cdot t}\right) \quad \textbf{[7]}$$

thus, the first order motor accumulation rate constant, $k_{acc}$, is:

$$k_{acc} = k_{on}^{2D} \cdot [Kin]_{tot} + k_{off}$$

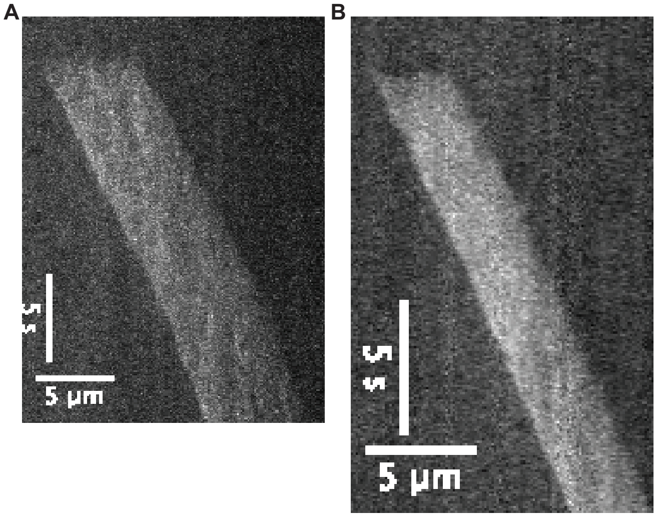


**Fig. S1.** Kymographs of motor accumulation at motor densities of ~40 /μm$^2$ (A) and ~120 /μm$^2$ (B) on membrane without cholesterol.

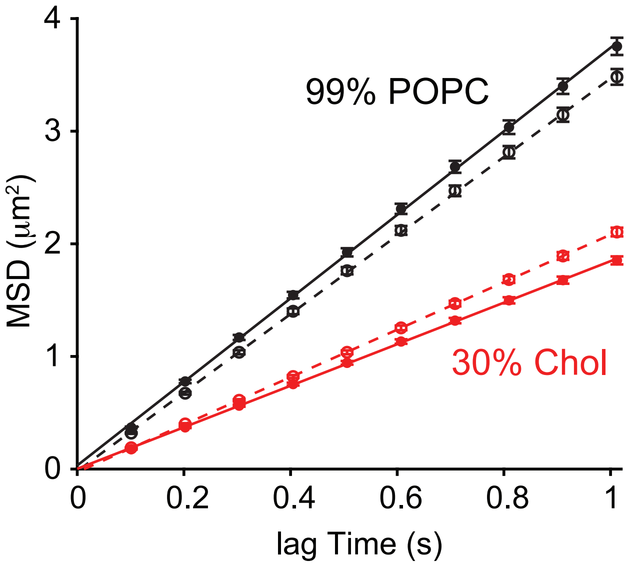


**Fig. S2.** Diffusivity of membrane-bound kinesin-1 motors by mean-squared displacement (MSD) analysis. Black closed circles and solid line show data and fit for bilayer composition of POPC:DSPE-PEG(2000)-Biotin:Atto 647N DOPE at 99.9:0.1:0.05. From fit, D = 0.93 ± 0.03 $\mu m^2/s$ (N = 193 tracked molecules, mean ± 95% confidence bound of fit). Black open circles and dashed line show data and fit for bilayer composition of POPC:DSPE-PEG(2000)-Biotin:Atto 647N DOPE at 99.4:0.6:0.05. From fit, D = 0.87 ± 0.02 $\mu m^2/s$ (N = 271). Red closed circles and solid line show data and fit for bilayer composition of POPC:cholesterol:DSPE-PEG(2000)-Biotin:Atto 647N DOPE at 69.9:30:0.1:0.05. From fit, D = 0.46 ± 0.01 $\mu m^2/s$ (N = 301). Red open circles and dashed line show data and fit for bilayer composition of POPC:cholesterol:DSPE-PEG(2000)-Biotin:Atto 647N DOPE at 69.4:30:0.6:0.05. From fit, D = 0.53 ± 0.01 $\mu m^2/s$ (N = 254). Error bars indicate SEM.

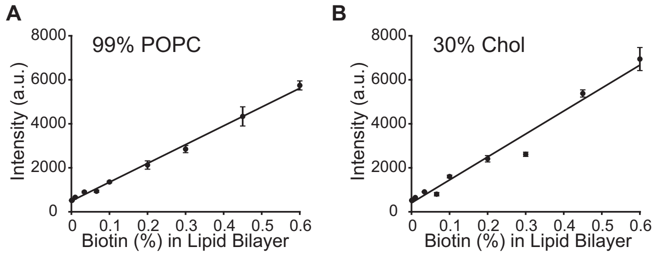


**Fig. S3.** Calibration curve of fluorescence intensity of membrane-bound K560GFPAviC against equivalent biotin concentration in lipid bilayer (DSPE-PEG(2000)-Biotin × spiking ratio). At high motor densities, saturating amounts of motors were added, so motor density was controlled by the fraction of DSPE-PEG(2000)-Biotin (0.1% - 0.6%) in the lipid bilayer. At low motor densities, sub-stoichiometric motors were added to membrane containing 0.1% DSPE-PEG(2000)-Biotin. Black closed circles show data averaged from 3 experimental days. Error bars indicate SEM. Lines are linear fits.

**Table S1.** Summary of diffusivity of lipids and membrane-bound kinesin-1 for different membrane compositions.

| Membrane composition (mol%) | | | Lipid diffusivity* | Motor diffusivity † |
|---|---|---|---|---|
| POPC | Cholesterol | DSPE-PEG(2000)-Biotin | $\mu m^2/s$ (FRAP) | $\mu m^2/s$ (MSD) |
| 99 | 0 | 0.1 | 1.02 ± 0.18 (N = 3) | 0.93 ± 0.03 (N = 193) |
| 99 | 0 | 0.6 | 0.88 ± 0.04 (N = 3) | 0.87 ± 0.02 (N = 271) |
| | | Average†† | 0.95 ± 0.14 (N = 6) | |
| 69 | 30 | 0.1 | 0.26 ± 0.04 (N = 3) | 0.46 ± 0.01 (N = 301) |
| 69 | 30 | 0.6 | 0.25 ± 0.01 (N = 3) | 0.53 ± 0.01 (N = 254) |
| | | Average†† | 0.25 ± 0.03 (N = 6) | |

* Values are mean ± SD.

† Values are mean ± 95% confidence bound of fit.

†† Combined data for membrane containing 0.1% DSPE-PEG(2000)-Biotin and 0.6% DSPE-PEG(2000)-Biotin.

**Movie S1.** Kinesin-1 accumulation on MT in 2 mM ATP (upper panel) or 2 mM AMPPNP (lower panel) at ~40 (left column), ~120 (center column) and ~250 (right column) motors/μm$^2$. Movie plays in real time.

**Movie S2.** Incorporation of cholesterol in POPC lipid bilayers reduces diffusivity of lipids. Movie shows FRAP for two lipid compositions: POPC:DSPE-PEG(2000)-Biotin:Atto 647N DOPE at 98.9:0.6:0.5 and POPC:cholesterol:DSPE-PEG(2000)-Biotin:Atto 647N DOPE at 68.9:30:0.6:0.5. Movie plays at 50 fps. Scale bar: 30 μm.
